# High-quality co-expression networks for accurate gene function predictions in the fungal cell factory *Aspergillus niger* and beyond

**DOI:** 10.1101/2023.07.28.550800

**Authors:** Paul Schäpe, Stephan Starke, Tabea Schuetze, Evelina Basenko, Sascha Jung, Timothy Cairns, Vera Meyer

## Abstract

Co-expression networks have recently emerged as a useful approach for updating and improving gene annotation at a near-genome level. This is based on the hypothesis that function can be inferred by delineating transcriptional networks in which a gene of interest is embedded. In this study, we generated a co-expression network for the filamentous cell factory *Aspergillus niger* from 128 RNA-seq experiments. We confirm that over 70% of the >14,000 *A. niger* genes are represented in this network and show that gene functions can be accurately predicted as evidenced by analysis of various control sub-networks. Our analyses further indicate that this RNA-seq co-expression network has a higher predictive power compared to the microarray co-expression network that we published in 2019. To demonstrate the potential of the new co-expression network to unveil complex and non-intuitive predictions for gene regulation phenomena, we provide here new insights into the temporal, spatial and metabolic expression profile that connects a secreted antifungal peptide with mycelial growth, asexual development, secondary metabolism and pectin degradation in *A. niger*. To empower biologists to generate or apply co-expression networks in the fungal kingdom and beyond, we also demonstrate that (i) high quality networks can be generated from only 32 transcriptional experiments; (ii) such low numbers of experiments can be safely compensated for by using higher thresholds for defining co-expression pairs; and (iii) a ‘safety in numbers’ rule applies, whereby experimental conditions have limited impacts on network content provided a certain number of experiments are included.

## Introduction

The fungal kingdom contains important model organisms, globally applied microbial cell factories, promising sources of renewable biomaterials, foodstuffs, and devastating pathogens of humans, plants, and other species (1, 2). As is the case for other kingdoms, fungal biologists have generated impressive compendiums of annotated genome sequences, which are integrated into publicly available databases for use by the research community (3–5).

However, most fungal genomes encode several thousand so called ‘hypothetical’ genes which lack any predicted function (6). Additionally, there are significant limitations in automated Gene Ontology (GO) annotation or by inferring function from reference systems such as *Saccharomyces cerevisiae*. Generating and interrogating co-expression networks has recently been established as a highly accurate and robust approach for predicting the function(s) for thousands of fungal genes (7–12). Such predictions using transcriptional networks are based on the principle that genes robustly co-expressed over multiple conditions are likely to function in the same, or similar, biological processes or metabolic pathways (7). Thus, by delineating co-expression networks for individual genes across diverse transcriptional experiments, function for a respective gene can be inferred by interrogating other genes in the network (7–12).

In 2019 we published a gene co-expression network for the industrial cell factory *Aspergillus niger* based on 155 microarray experiments (7). This resource was successfully used to predict six putative transcription factors (MjkA – MjkF) that function as global or pathway-specific regulators in secondary metabolism (7, 12). These genes were previously uncharacterized in the lab yet are deeply embedded in secondary metabolite transcriptional networks. We demonstrated that over-expression of *mjkA*-*F* using a Tet-on synthetic gene switch that allows loss-of-function and tuneable gain-of-function experiments can be used to concomitantly activate multiple secondary metabolite pathways (7, 12). We furthermore showed that exploration of this co-expression resource can unravel unexpected regulators that modulate production of secreted proteins and organic acids (9), which are the two main classes of industrially relevant products of *A. niger* (6). In fact, sub-networks predicted that the citric acid cycle and the protein secretion machinery are transcriptionally coupled via the Golgi organelle, a hypothesis which could not be derived from other approaches for gene functional prediction (e.g., GO/KEGG annotation, orthology with informant genomes). We could thus experimentally verify using the Tet-on switch that genes predicted to encode Golgi-localized activators or deactivators of the GTPase ArfA (*ageB*, *secG*, and *geaB*) are required for mitochondrially-localized citric acid production in *A. niger* (9).

Co-expression resources were also used by other groups for the human-infecting dimorphic yeast, *Candida albicans*, to accurately predict that *ccj1* is involved in cell cycle progress (10), although this gene lacked any functional predictions (e.g., GO terms), and had dubious orthology with other yeasts, e.g., *Saccharomyces cerevisiae*. More recently, co-expression networks were integrated with functional genomic resources (a *C. albicans* Tet-on driven conditional expression strain library) to predict and confirm the functions of three uncharacterized essential genes, which demonstrated that the *gln4* gene product is the target of the antifungal compound N-pyrimidinyl-β-thiophenylacrylamide (11). Thus, co-expression network construction and interrogation has proven to be a highly useful approach for accurate predictions of gene function from industrial and pathogenic fungi.

Nevertheless, several challenges and outstanding questions remain for the widespread applications of co-expression networks in the fungal kingdom and beyond. Do RNA-seq and microarray technologies perform comparably for network construction? How many transcriptional samples should be included for network construction? What are useful thresholds for defining robust co-expression across a given dataset? Can a limited amount of input data (i.e., low number of experimental conditions) be compensated for by more robust thresholds for defining co-expressed genes (i.e., high correlation coefficients between genes)? Are there specific experimental conditions that should be included to maximize the utility of the network (e.g., various nitrogen or carbon sources)? These questions are especially important for the many hundreds of fungal species that are of major interest yet lack a well-established research community, such as emergent pathogens (13), or promising new strains/species identified during bioprospecting efforts (14). Co-expression networks for such species will likely start from scratch, cannot rely on mining publicly available datasets, and therefore must effectively utilize time and cost intensive resources.

In this study, we address the above questions using the cell factory *A. niger*. Multiple RNA-seq networks consisting of over 100 million co-expression pairs were generated from 128 biological experiments and compared to the existing genome-wide network derived from 155 microarray experiments. We expand on the previously described pseudo random network generation approach (7) to demonstrate that biologically meaningful co-expression networks can be derived from only 32 transcriptional experiments. We also generate guideline Spearman correlation coefficients for various network construction scenarios. By interrogating positive control networks of genes with verified functions in *A. niger*, we demonstrate that most networks consist of a core of robustly co-expressed genes, and a subset of context dependent expression relationships that vary with input data. Finally, we generate a novel hypothesis for the function of the antifungal protein AnAFP and confirm this by generating a Tet-on conditional expression mutant. Taken together, our study provides a co-expression resource which updates our previous genome annotation of *A. niger* (7) and a set of guidelines that can be used to build co-expression networks for any organism in the fungal kingdom and beyond.

## Materials and methods

### Network construction

#### Data mining

During January 2022 the NCBI Sequence Read Archive (SRA - https://www.ncbi.nlm.nih.gov/sra) was searched for “*Aspergillus niger* CBS 513.88”. 381 publicly available transcriptome datasets for the strain CBS 513.88 were found. Additionally, 85 unpublished samples for CBS513.88 were added from our group leading to a total of 466 RNA seq datasets. Supplemental File 1 lists all datasets and relevant information regarding these files. Some samples were excluded due to insufficient information on the cultivation conditions deposited at NCBI-SRA.

#### Data processing and quality control

SRA-Toolkit version 2.10.0 was used for downloading and unpacking the datasets. Paired read data sets were merged using BBMap_37.92. FastQC 0.11.9 was used to investigate the overall data quality. Quality control was done in an automated fashion and all generated FastQC report files were manually curated. Several samples could not be downloaded correctly or did not meet the required level of quality and had to be excluded from further analysis. Finally, 348 datasets reflecting 128 different cultivation conditions out of the initial 466 samples were used for network creation. The detailed overview of the datasets is listed in Supplemental File 1.

#### Mapping and normalizing raw counts

For Mapping the tool STAR (15) was used with suggested standard parameters. Reads were mapped against the Genome of strain CBS513.88 version 2 accessed in February 2020 (https://www.ncbi.nlm.nih.gov/genome/429?genome_assembly_id=54477). The read counts were normalized to Transcripts per Million reads (TPM) by a private python script. All TPM values for all genes over all conditions are given in Supplemental File 2.

#### Spearman calculation and constructing the main network

To avoid unfounded statements of biological relations of low expressed or transcriptionally silent ORFs, an expression variance threshold was set to at least 10% of the mean variance of all genes over all conditions. Any ORF with a lower variance in its expression in the 128-condition data set was not considered for any of the Spearman coefficient calculations. 4,234 out of the 14,422 predicted ORFs had a variance lower than this threshold. For network creation for each pair of ORFs/gene the Spearman’s rank coefficient (16) was calculated using Python 3.17 with the package SciPy version 1.7.1 (17). The calculated Spearman coefficients can be interpreted as the connections/edges describing the interaction of the genes/nodes in the network.

#### Creating test networks using a range of RNA-seq input scenarios

Co-expression networks derived from 8, 16, 32, 64, or 96 RNA-seq experiments that were randomly selected from the full 128 dataset were generated as described above. We termed these datasets ‘test’ networks, which were used to confirm the minimal amount of RNA-seq experiments to generate high-quality co-expression resources. Technical replicates were generated for test networks, with a minimum of 3 (8 and 16) and maximum of 6 (32, 64, 96) randomly selected RNA-seq experiments. We also generated two networks from RNA-seq input that were intentionally biased, specifically experiments that were carbon rich or lacking nitrogen. RNA-seq samples used for respective network construction are given in Supplemental File 3.

#### Construction of pseudo random co-expression networks

Pseudo random co-expression datasets were created as described previously (7). In this approach, gene expression values were randomized prior to network construction so that gene transcript values (TPM) at a given condition were randomly re-assigned to another condition (7). Pseudo randomized datasets were generated using RNA-seq samples described for the main coexpression resource (n = 128) or test networks (n = 8, 16, 32, 64, 96, Supplemental File 3). We calculated the frequency of Spearman correlation coefficient *n* in the pseudo random networks vs corresponding test networks (i.e., generated from either 128, 96, 64, 32, 16 or 8 RNA-seq samples). We accepted that a 95% frequency of Spearman *n* in the main/test vs corresponding pseudo randomized network was an acceptable threshold.

#### Analysis of co-expression networks

Various key features of the different networks (number of correlations/ genes included in network/ transcription factors included in network/ etc.) were calculated while creating these networks. The classification of genes into transcription factors, secondary metabolite core genes and hypothetical genes were obtained from FungiDB (7). Individual gene sub-networks were created as previously described (7), whereby for gene *n*, all co-expression pairs above a given correlation threshold were reported. Gene ontology enrichment analysis (GOE) was performed by using the FungiDB GEO tool. Network and sub-network visualization was done with Cytoscape 3.9.1 (18). Heatmaps were created by using Python and MatPlotLib 9.1 and Seborn 9.2.

### Molecular cultivation approaches

#### Strains generated in this study

Plasmid construction was performed via Gibson assembly. Transformation of *A. niger* was conducted as described earlier (19). Single copy integration of the exogenous cassette in the recipient genome was validated via Southern analysis. For STS4.10, the *anafp* (1kb upstream of the ORF) was substituted via double homologous recombination with the Tet-On promotor resulting in PfraA-rtTA2-TtrpC-tetO7-PmingdpA--anafp-Tanafp. Selection was performed via the *pyrG* gene from *A. oryzae*. VG8.27 originated from Meyer et. al. (20).

#### Bioreactor cultivations

For RNA-seq data for *A. niger* cultivated under low nitrogen and steady-state conditions (Supplemental File 1), two BioFlo 310 bioreactors (New Brunswick Scientific, Edison, USA) were run with 5 kg cultivation medium with low nitrogen (per litre: 29.7g glc·H_2_O, 0.8 g NH_4_Cl, 1.5g KH_2_PO_4_, 0.5 g KCl, 0.5 g MgSO_4_·7H_2_O and 1 ml of 1000x concentrated trace element solution) and inoculated with 1.0 · 10^6^ spores/ml. Temperature of 30°C and pH 3 were kept constant, the latter by computer controlled addition of 2 M NaOH or 1 M HCl, respectively. The sterile air flow was 1 L/min. Dissolved oxygen tension (DOT) and pH were measured with autoclavable sensors (Mettler Toledo). Online measurements for O_2_ and CO_2_ off-gas were done by an EX-2000 analyzer (New Brunswick Scientific, Edison, USA). Dissolved oxygen tension (DOT) and pH were measured with autoclavable sensors (Mettler Toledo). The dilution rate was set to 0.1 h^-1^ after 2.5 g/kg biomass wer reached. During steady state (after the 5^th^ residence time), samples for RNA extraction were taken and the medium was changed to glucose limiting minimal medium (per litre: 8.0 g glc·H_2_O, 1.5 g NH_4_Cl, 1.5g KH_2_PO_4_, 0.5 g KCl, 0.5 g MgSO_4_·7H_2_O and 1 ml of 1000x concentrated trace element solution), also here samples for RNA extraction were taken during steady state (8th residence time of the glucose-limiting medium).

Strain STS4.10 and VG8.27 (20) were cultivated in a 5L bioreactor as described in Jørgensen et al.(21). Complete medium was used (MM + 1% yeast extract and 0.5% casamino acids, without doxycycline). Hyphae were harvested in the mid exponential growth phase by filtration of 7 ml through a nitrocellulose membrane. The biomass on the filter was transferred to agar plates containing minimal medium without glucose in the presence or absence of 5 µg/ml doxycycline. A second nitrocellulose membrane was placed on top to repress sporulation (22). Cultures were subsequently starved for 35 hours and ∼300 mg were used for RNA extraction. RNA quality was checked by capillary electrophoresis by the sequencing company.

### Differential gene expression analysis

Paired end sequencing was performed using Illumina nova seq with a minimum of 10 million reads per sample. Sequencing quality was checked using multiQC 6, untrimmed reads were mapped against the genome of *A. niger* N402 (ATCC64974) using STAR 7. Differential gene expression was conducted using DESeq 2. Resulting genes were filtered for p-values lower than 0.05 (FDR corrected, Benjamini-Hochberg) and a log2 Fold-change higher than 1.5. Genes from ATCC64974 were translated into syntenic orthologs of CBS513.88 and GO enrichment was conducted with Bonferroni corrections and Fisher’s exact test by the use of FungiDB (3).

### Luciferase assay

100 µl of 25 mM Luciferin solution was added to a cooled 20 ml CM agar plate. 10^4^ spores of PK2.9 (Panafp, *luciferase*, Δ*anafp*) were then spotted in the center of the plate. The plate was incubated at 30°C for a period of 3 days. At 18, 30, and 60 hours, photographs were taken under both normal light conditions and complete darkness using a Canon 6D DSLR camera with a 50mm lens set at f1.4 and an exposure time of 30 minutes and an ISO of 3200. Finally, the dark frame was post-processed in Lightroom for noise reduction and tonal correction, and then overlayed with the light frame (converted to monochrome) for analysis.

### Growth assays

The protocol used in this study is based on the method previously described (23). 10^4^ spores of STS4.10 (Tet-on *anafp*) and VG8.27 (Tet-on *luciferase*) were added to the center of minimal medium agar containing 1% glucose, and incubated at either 30°C or 42°C. Alternatively, 1% pectin was used as a substrate (with glucose omitted) and incubated at 30°C. The plates were incubated for 72 hours, and the colony diameter was measured using ImageJ in three independent replicates. This data was used to calculate growth coefficients.

## Results

### Generating an *A. niger* co-expression network based on RNA-seq data

For co-expression network construction, we retrieved RNA-seq data for the *A. niger* wildtype strain N402 and its derivative isolates that were publicly available or from our own labs (Supplementary File 1). These data include diverse cultivation conditions, including solid and submerged growth during bioreactor or shake flask cultivations on different carbon (e.g. pentoses, hexoses, xylan, straw) and nitrogen sources (e.g. amino acids, nitrate, ammonium), different developmental stages (e.g. germination, mycelial growth, starvation, conidiophore development and sporulation), a wide range of mutant isolates affected in stress resistance, protein secretion and secondary metabolism, and data from co-cultivations with other fungi on lignocellulosic biomass. Notably, these data and the previous microarray derived co-expression network (7) lacked growth under nitrogen limitation. As nitrogen limitation is known to affect secondary metabolism in filamentous fungi (24, 25), we added this condition by running a chemostat cultivation of the wildtype strain N402 using ammonium as the limiting nitrogen source (see Materials and Methods). Thus, a total of 128 unique *A. niger* transcriptional experiments were used to generate the co-expression resource (Supplementary File 1). Co-expression values were calculated based on Spearman’s rank correlation coefficients (16) and preliminary visualisation and interrogation of the network was conducted with coefficients above our previously defined stringent threshold of > +/-0.5 (Figure 1A, (7)). This demonstrated 70% coverage of predicted *A. niger* genes in the network (n = 9,952, >350,000 co-expression pairs) and included functional categories that are of major research interest. This includes, for example, 70% of putative *A. niger* transcription factor encoding genes (n = 287, Table 1). Additionally, we found that 48% of hypothetical genes (n = 1,589, Table 1) were included in this network. We also interrogated expression values for the 79 predicted biosynthetic gene clusters (BGCs, (26)) across the 128 conditions, confirming that most were expressed in one or more condition (Figure 1B), although these genes are conventionally assumed to be transcriptionally silent during laboratory culture (25). 38 core genes (e.g. polyketide synthases or non-ribosomal peptide synthases) of these BGCs were found to belong to sub-networks, confirming that even rare/low expressed genes are included in the dataset. For clarity, we refer to this dataset as the ‘main network’ throughout the remainder of this study.

**Figure 1:**
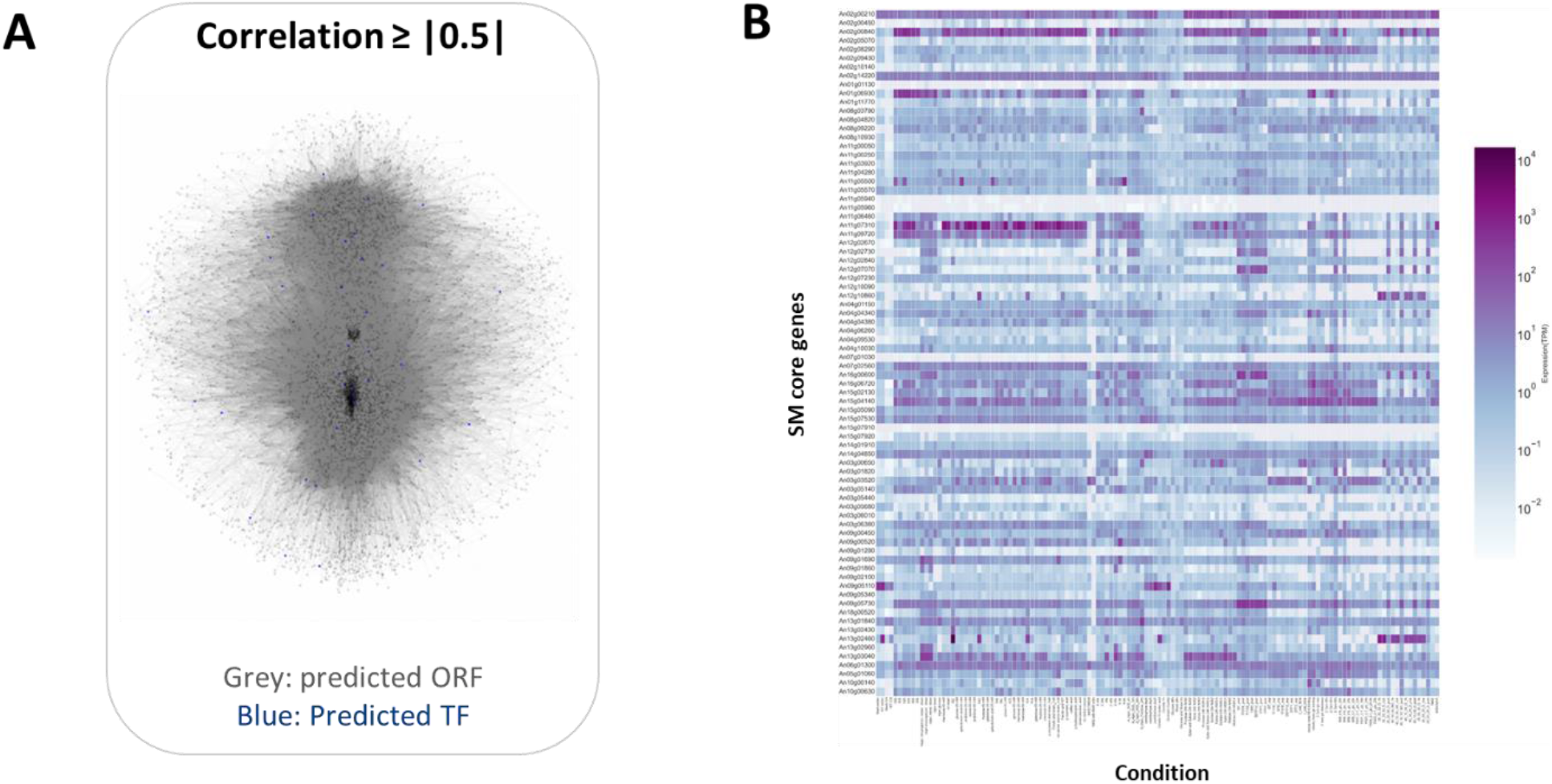
(A) Schematic representation of the co-expression network generated in this study. Genes are represented by boxes, and connections by lines. The 287 transcription factor encoding genes are highlighted in blue. **(B)** Normalised expression values for 81 core genes of 79 secondary metabolite biosynthetic gene clusters across the 128 RNA-seq experiments. Most genes are expressed in one or more conditions.

**Table 1:** Key features of networks generated in this study. *Using minimal correlation coefficients (Table 2), **Average values from 6 networks, ***Spearman 0.5/-0.5, 280 conditions

|  | RNA-seq* |  |  |  |  |  | Microarray network***<br>(7) |
| --- | --- | --- | --- | --- | --- | --- | --- |
|  | Main Network<br>(128) | Network<sup>96**</sup> | Network<sup>64**</sup> | Network<sup>32**</sup> | Main+N<br>(124) | 64Carbon |  |
| Total number of genes in the network (out of 14,422 genes in total) | 10,188 | 10,188 | 10,177 | 10,151 | 9,796 | 10,184 | 9,579 genes |
| Total number of connections | Pos:<br>5,582,576<br>Neg:<br>517,253 | Pos:<br>3,638,021<br>Neg:<br>195,513 | Pos:<br>2,539,031<br>Neg:<br>66,367 | Pos:<br>1,362,460<br>Neg:<br>/// | Pos:<br>1,110,812<br>Neg:<br>21,838 | Pos:<br>2,798,765<br>Neg:<br>128,132 | Positive n =<br>5,476,855<br>negative n =<br>3,613,374 |
| Predicted transcription factor encoding genes in the total network (out of total 409 in total) | 287 | 287 | 287 | 283 | 280 | 287 | 267 |
| Predicted hypothetical genes in the network (out of 3,302 in total) | 1,589 | 1,589 | 1,587 | 1,585 | 1,525 | 1,589 | 1,243 |
| Predicted secondary metabolite core genes (out of 79 BGCs) | 38 | 38 | 38 | 38 | 38 | 38 | 49 |

### Defining minimal correlation thresholds and data input

In order to determine minimal amounts of experimental input for meaningful network construction, we generated co-expression networks derived from 8, 16, 32, 64, or 96 RNA-seq experiments which were randomly selected from the full compendium of 128 conditions (Figure 2). These networks, which we named ‘test networks’, model various data availability scenarios that may be encountered when generating a co-expression resource, e.g., very limited (n = 8, 16), moderate (n = 32, 64) to high (n = 96) availability of transcriptional experiments.

**Figure 2:**
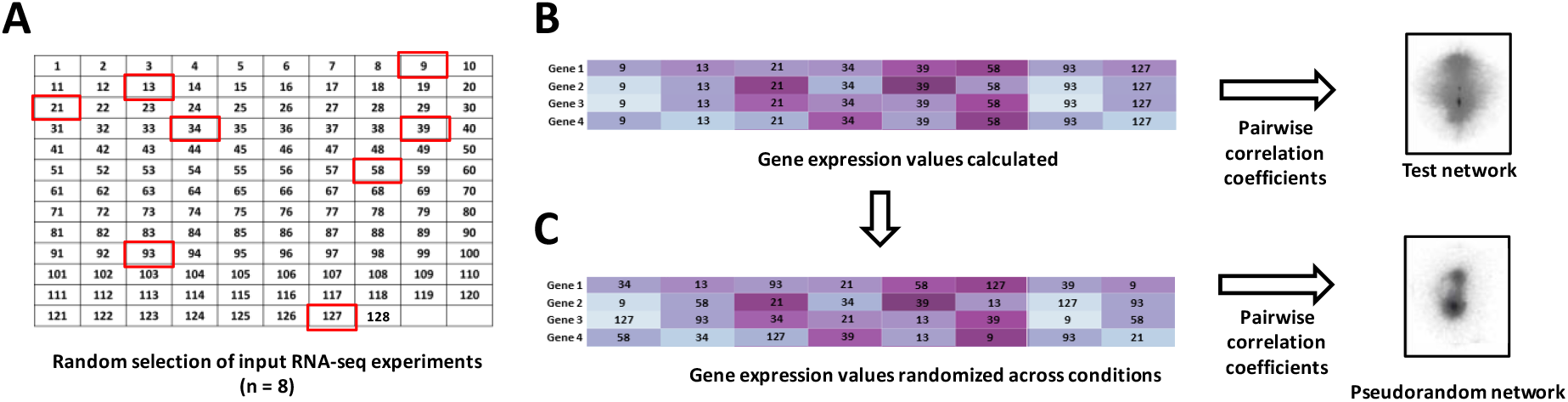
Schematic representation of network generation and pseudo randomisation. Generation of a network using 8 randomly selected RNA-seq experiments from the database of 128 is depicted (A). Expression values are normalised for *A. niger* genes (4 exemplars shown in B). Note, colour intensity indicates expression level, while numbers indicate the RNA experiment from which this value was derived. Based on these data, pairwise Spearman correlation coefficients are calculated between every gene in the *A. niger* genome, to generate a test network. For pseudo randomisation (C), normalised gene expression values are randomised across experiments. Thus, the distribution of gene expression values is identical between test and pseudo random network (note identical colour intensity between B and C), yet expression of gene *n* in experimental condition *x* is biologically incorrect. From these data, pairwise correlation coefficients are recalculated between every gene pair. In the pseudo random network (C), correlation between gene pairs can only occur due to chance, whereas in the test network (B), gene correlation can occur due to biologically based co-expression.

For the main network (Figure 1A) and each test network, we then applied a pseudo random approach as described earlier (7), whereby normalised expression values for each gene were randomised across experimental conditions, after which Spearman correlation coefficients were recalculated (Figure 2B and C). This generated pseudo randomized networks that (i) have identical distribution of gene expression values compared to the test network control, yet (ii) are biologically meaningless, i.e., the degree of gene co-expression will occur due to chance rather than biological phenomena. Thus, pairs of biological (test) and nonsense (pseudo randomized) networks were generated from 8, 16, 32, 64, 96 or 128 RNA-seq experiments.

In order to determine minimal Spearman correlation cut-offs, we calculated how frequently defined Spearman correlation coefficients were found in the six test networks (8, 16, 32, 64, 96, 128 RNA-seq experiments) vs the six pseudo randomized networks at each of the data availability scenarios (Figure 3). We considered that a 95% frequency of a given Spearman value occurring in the biological datasets when compared to the pseudo random datasets was a reasonable threshold for determining minimum correlation coefficients (Figure 3, Table 2, and Supplemental File 4). As expected, this minimum Spearman correlation coefficient threshold was inversely correlated with the number of RNA-seq experiments used to build the network (Figure 3). We found that this threshold was never reached when only 8 or 16 RNA-seq datasets were used, and consequently we consider that these numbers of experiments are insufficient for building co-expression resources. Our approach demonstrates that 32 networks are appropriate for network construction, provided that a reasonably high correlation threshold is used (0.56). This data further suggest that our previously used Spearman correlation coefficient threshold of 0.5 for the microarray-based network (7) was likely unnecessarily stringent, and probably resulted in high rates of false-negative results (i.e., excluding correlations from gene sub-networks that were informative for gene function). Notably, correlation thresholds differed for positive and negative Spearman thresholds (Table 2), a phenomenon that we discuss further below.

**Figure 3:**
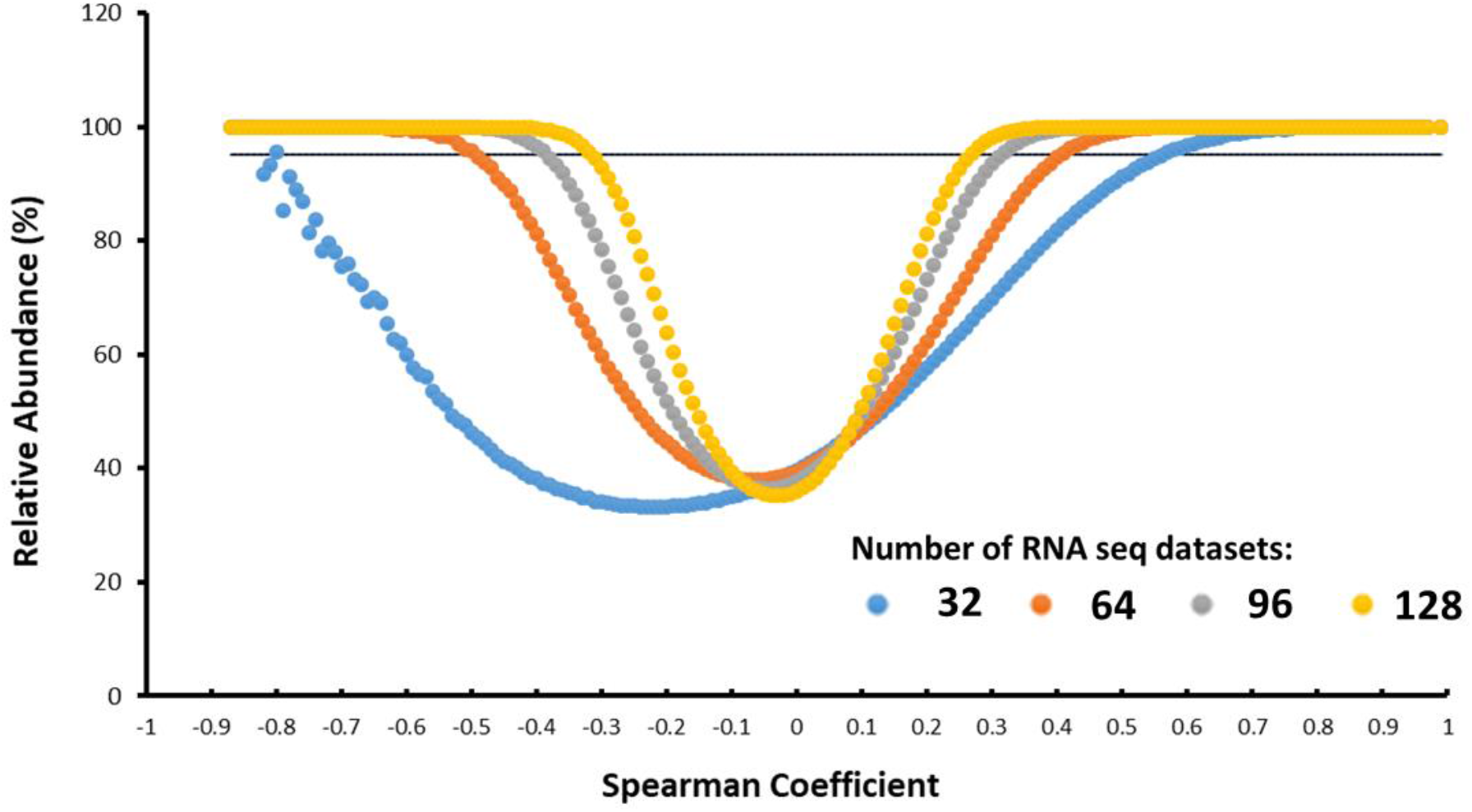
Frequency of Spearman correlation coefficients in biological or pseudo randomised network pairs. Spearman correlation coefficients between gene pairs were calculated to two decimal places and counted in either biological (main and test) or pseudo random networks, with data reported as % abundance in main and test networks. As expected, higher Spearman correlation coefficients are more frequently found in the biological networks. For example, a correlation coefficient >0.8 is never observed from >400 million pseudo random gene pairs, and consequently correlation coefficients above this value must be co-expressed due to biological phenomena. Note that the number of RNA-seq datasets from which the network was constructed impacts the frequency of Spearman correlation coefficients in biological vs pseudo randomised networks. For example, a Spearman of 0.3 occurred about 30% of the time in randomised datasets when 32 RNA-seq samples were used for network construction. In contrast, this value was never observed due to chance when 128 networks were used. We considered 95% frequency in biological vs pseudo randomised data to be a cut off for our minimum advised Spearman correlation coefficients (depicted by a horizontal black line).

**Table 2:** Minimum Spearman correlation coefficients for biological networks derived from different numbers of RNA-seq datasets. Increasing Spearman cut-offs could be used to compensate for networks generated from lower numbers of experimental conditions. Advised minimum cut-offs indicate that values above this threshold give >95% confidence of biologically valid co-expression.

| Number of RNA-seq experiments | Advised minimum Spearman correlation cut-off for positive co-expression | Advised minimum Spearman correlation cut-off for negative co-expression |
| --- | --- | --- |
| 128 | 0.25 | -0.31 |
| 96 | 0.31 | -0.38 |
| 64 | 0.40 | -0.48 |
| 32 | 0.56 | -0.90 |
| 16 | None | None |
| 8 | None | None |

### Predicting gene biological functions from the main co-expression network

To test the validity of the main network and our estimated minimum coefficient thresholds (correlation coefficient >0.25, Table 2), we interrogated sub-networks for genes where biological function has been comprehensively determined and which are of our research interest. This includes genes encoding (i) the secreted glucoamylase, GlaA, which is used for the industrial saccharification of starch to glucose (27), (ii) the nonribosomal peptide synthetase SidD, which is crucial for adaptation to low iron environments through biosynthesis of the siderophore triacetylfusarinine C (TAFC, (28)), and (iii) the secreted peptide AnAFP, which is a potential lead compound for antifungal drug development due to its fungal-specific mode of action (29).

Analysis of control sub-networks at both gene ontology (GO) and gene-level revealed clear enrichment of many expected biological processes, subcellular parts, metabolic pathways, and membrane transporters. As depicted and summarized in Figures 4, 5 and Table 3, the *glaA* gene was co-expressed with 461 genes among which were genes encoding other starch degrading enzymes (e.g., the alpha-amylase *amyA* (An04g06930)) and the *glaA* sub-network was enriched with multiple processes important for the classical protein secretion route (30). The *sidD* sub-network contained 556 genes, was enriched for TAFC biosynthetic GO terms (GO:1900551, p = 0.04) and thus included strong co-expression of genes that reside in the contiguous TAFC biosynthetic cluster (An03g03560, An03g03620 (*ste6*), An03g03540 (*sidF*), An03g03550 (*sidH*), An03g03530 (*sidJ*)). Notably, the *sidD* sub-network also contained genes outside its predicted BGC that are nevertheless required for TAFC synthesis or predicted plasma membrane transport (An06g01320 (*sidI*) and An07g06240 (*mirD*), respectively (31). The *anafp* co-expression network harboured 2,651 genes and was enriched with 94 secreted proteins and enzymes (GO GO:0005576, p = 0.01), and a functional category equating to a predicted cell wall-related mode of action for this antifungal peptide (GO:0044347, p < 0.01). We have previously hypothesised that activation of autophagy may be an important component of antifungal activity of AnAFP (8). Indeed, clear co-expression relationships between *anafp* and nine predicted autophagy-associated genes (*atg9, atg11, atg12, atg15, atg16, atg18, atg22, atg24, atg29*; Figure 4) were found, several of which were among the strongest co-expressed pairs in the network (e.g., the predicted vacuolar lipase encoding gene *atg15*, Spearman 0.58). By plotting Spearman correlation values as a function of proximity, we could clearly delineate clustering of related processes within these three sub-networks, for example cell wall catabolism and autophagy in the *anafp* network (Figure 5). Taken together, analysis of these control networks confirms useful prediction of gene functions can be derived from the main network using our lowest advised minimum Spearman correlation coefficient.

**Figure 4:**
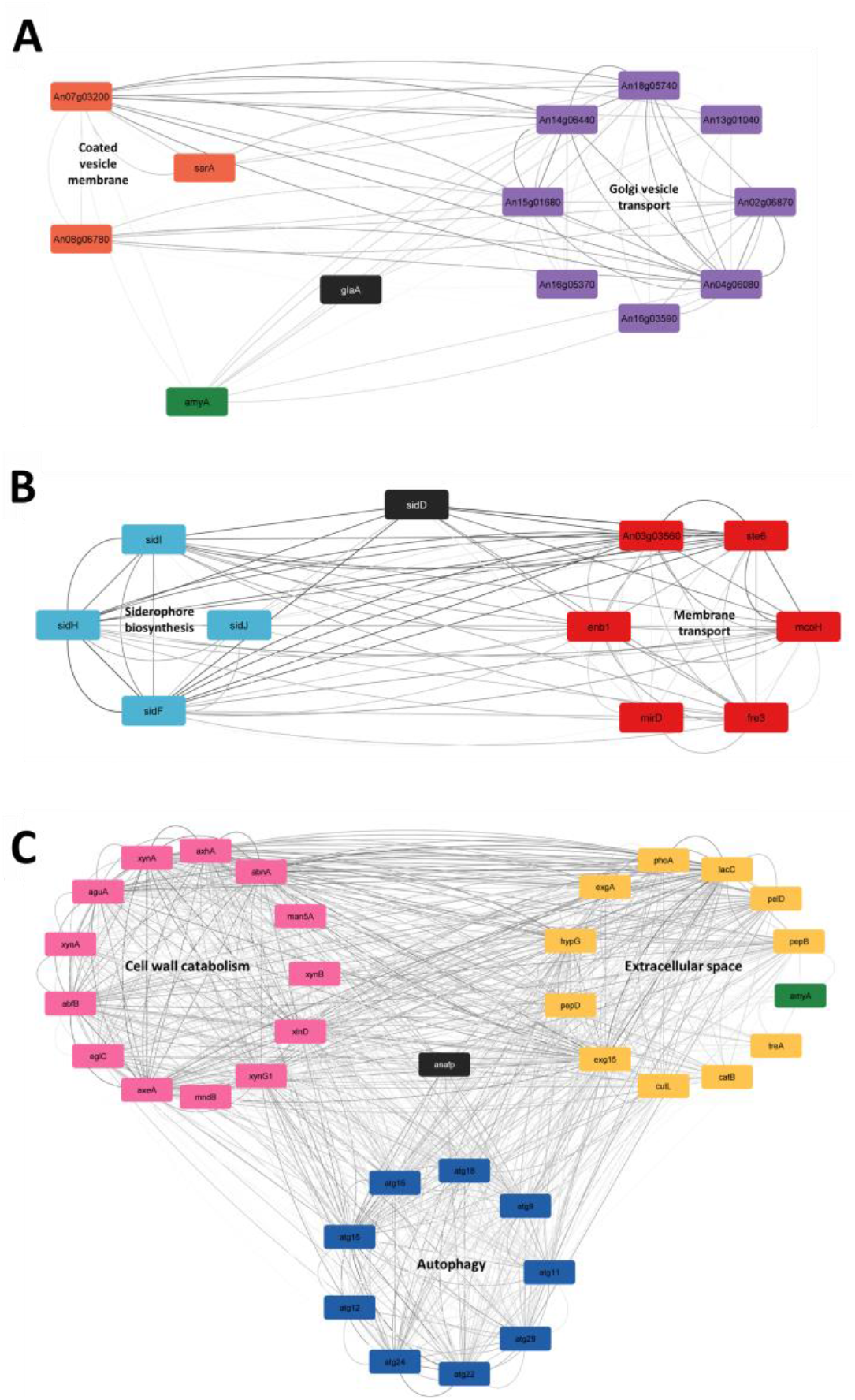
Schematic depiction of *glaA, sidD*, and *anafp* co-expression sub-networks. Genes are depicted in squares, with co-expression depicted by grey lines. Sub-network central genes are given in black boxes. Co-expressed genes with the sub-network central gene are arranged into different functional categories.

**Figure 5:**
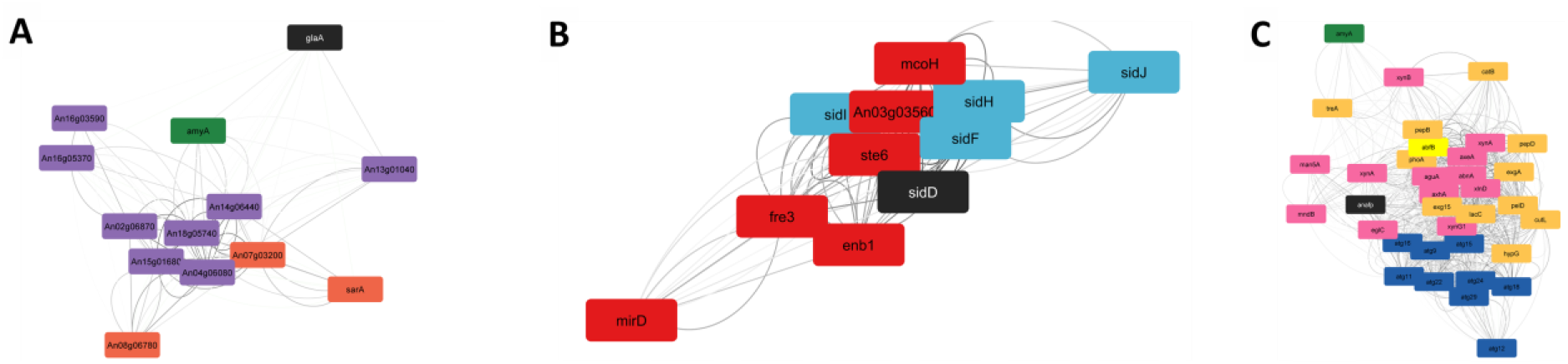
Schematic depiction of *glaA, sidD*, and *anafp* co-expression sub-networks with Spearman correlation coefficients plotted as a function of proximity. Genes are depicted in squares, with co-expression depicted by grey lines. Sub-network central genes are given in black boxes. Gene colours denote grouping into functional categories in Figure 4.

**Table 3.**
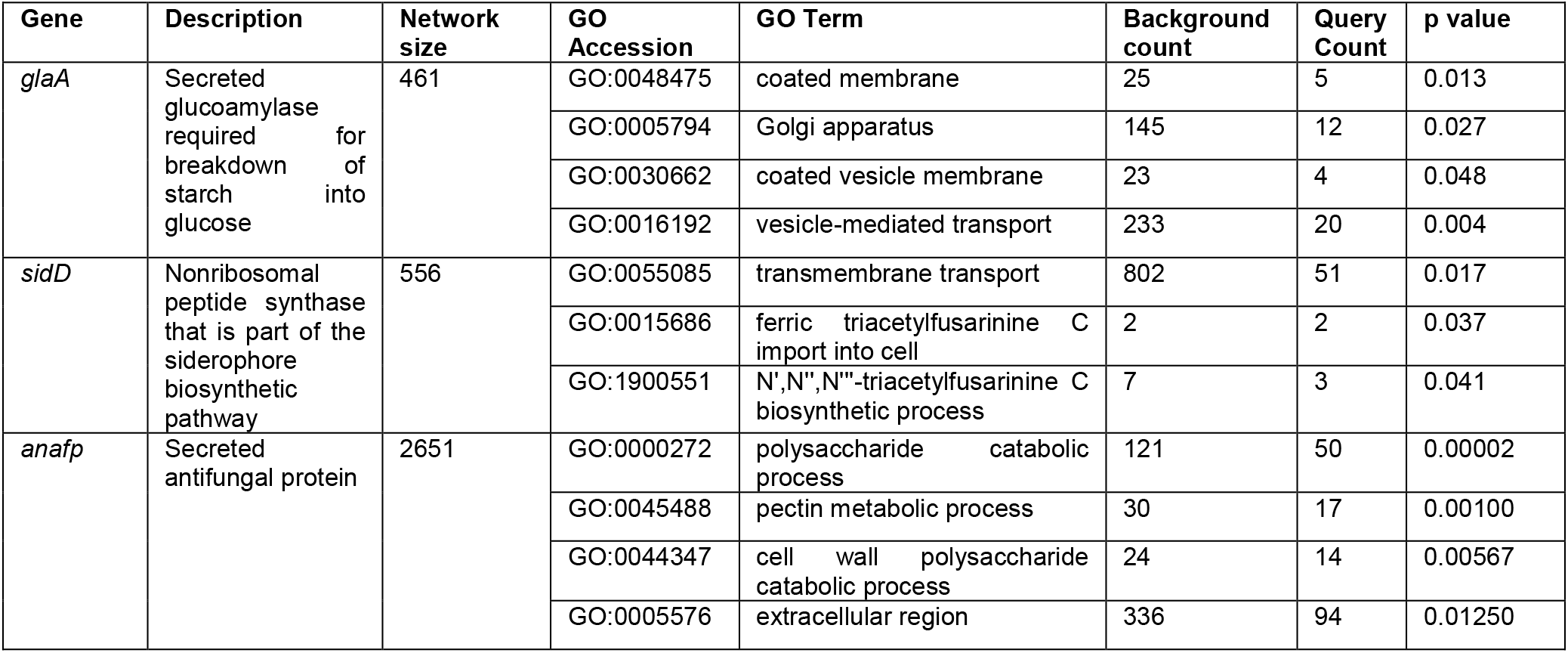
Selection of enriched GO-terms amongst *glaA, sidD* and *anafp* co-expression sub-networks.

### Comparing gene correlation coefficients between different RNA-seq networks

Next, we assessed how variation in *A. niger* cultivation conditions used for RNA-seq network construction impacted correlation coefficients and gene sub-network content. We therefore generated additional co-expression networks (n = 6) from either 32 or 64 randomly selected RNA-seq experiments and interrogated *glaA, sidD*, and *anafp* sub-networks from these datasets. Correlation coefficients for the top 30 genes from 6 sub-networks are visualised in Figure 6 (derived from 64 RNA-seq samples) and Supplemental Figure 1 (derived from 32 RNA-seq experiments). These demonstrate robust gene correlation coefficients across sub-networks/datasets, especially for the *glaA* sub-network (Figure 6A). Strikingly, all six *sidD* sub-networks showed a core set of five genes that are reproducibly the highest correlated with *sidD* (Figure 6B and Supplemental Figure 1), which either were part of the contiguous gene cluster (e.g., *sidF*, *sidH*), key TAFC biosynthetic genes (*sidI*), or predicted parts of the *A. niger* iron limitation response (putative multicopper oxidase encoding gene *mcoH*, An01g08960). Genes with *sidD* correlation coefficients below this core set were also found to be highly comparable across the six datasets, and in virtually every instance passed the lower correlation threshold for inclusion in the network. Interestingly, the largest deviation in sub-network content in this experiment was observed for *anafp* (Figure 6C), with two populations of sub-networks distinguishable based on individual gene correlation coefficients (sub-networks 1, 2 and sub-networks 3, 4, 5, and 6, Figure 6C). Clearly, these correlation coefficients vary based on the RNA-seq datasets from which the networks were built. Nevertheless, virtually all the depicted genes in Figure 6C pass the minimum correlation coefficient of 0.40 (Table 1), demonstrating that these genes are robustly co-expressed with *anafp* in all six test networks.

**Figure 6:**
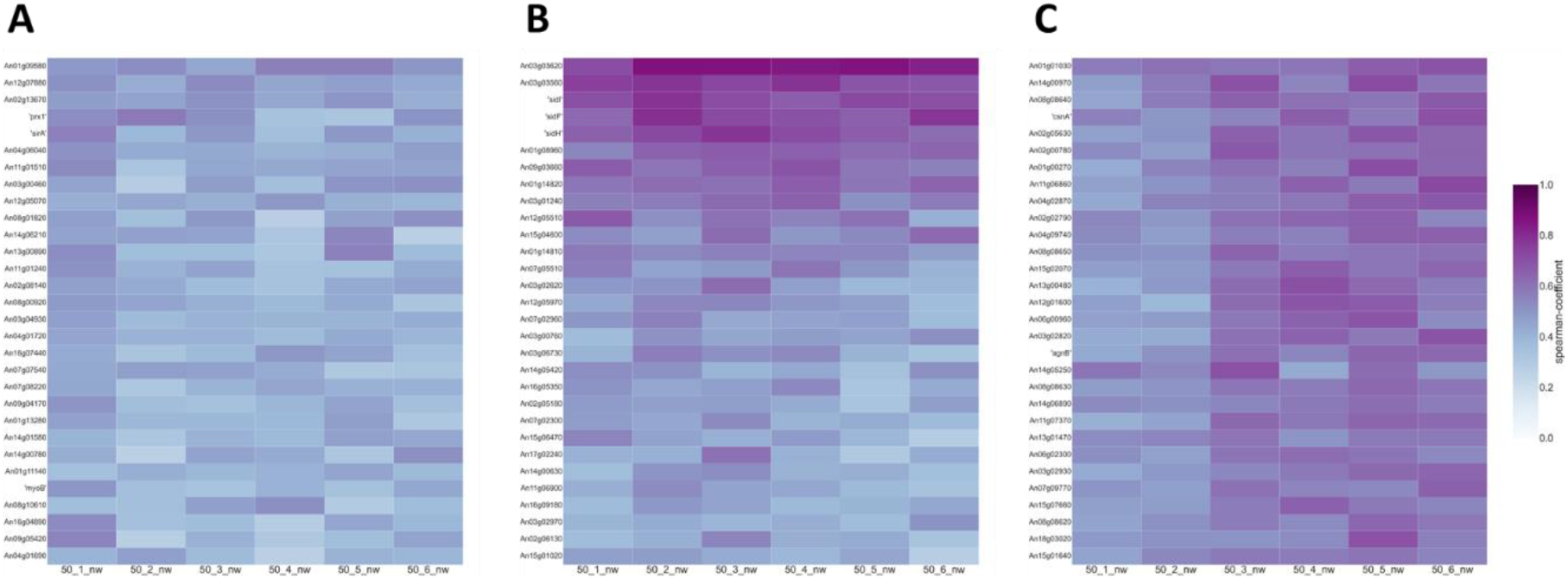
Correlation coefficients for *glaA, sidD*, and *anafp* across six different test networks derived from 64 experimental conditions that were randomly chosen. Top 30 highest co-expressed genes are depicted for either *glaA* (A), *sidD* (B), or *anafp* (C). Each column represents a distinct network derived from 64 randomly selected RNA-seq experiments. Colour intensity correlates with Spearman correlation coefficient.

Next, we tested how omission or enrichment of certain cultivation conditions might impact correlation coefficients in a network. Given the crucial role of nitrogen availability in cell growth, we hypothesised that a network derived from transcriptional experiments lacking nitrogen limitation might be different to the main network generated in this study. We therefore omitted the four nitrogen limitation RNA-seq experiments (Supplemental File 3), and calculated correlation coefficients from the remaining 124 datasets to generate what we named the ‘+N’ network. Interestingly, the *glaA, sidD,* and *anafp* sub-networks from the ‘+N’ network (correlation coefficient >0.25, Table 2) were virtually identical to those derived from the main network (Supplemental File 5). Thus, we conclude that the inclusion of nitrogen limited growth does not drastically impact sub-network content provided sufficient numbers of experiments are included (e.g., 124 RNA seq samples).

In order to test how cultivation in a range of carbon sources impacted gene coexpression, we generated a network derived from 64 carbon diverse RNA-seq experiments, which we termed the ‘64Carbon’ network (Supplemental File 3). When compared to a test network generated from 64 randomly selected samples (correlation coefficient >0.40, Table 2), the *glaA, sidD,* and *anafp* sub-networks were strongly impacted, both in the number of genes (nodes) and correlation coefficients (edges, Supplemental File 5). Despite these differences, accurate predictions of gene function were found in the 64Carbon derived sub-networks for both *sidD* (TAFC biosynthetic genes, GO:1900551, p < 0.002) and *glaA* (e.g., Golgi vesicle transport GO:0048193, p >0.001). Only the *anafp* 64Carbon subnetwork did not return enriched GO terms consistent with the function of the encoded protein, an observation that (i) demonstrates some networks are reduced in functional connectivity due to RNA-seq inputs and (ii) is consistent with the role of this protein in carbon/pectin catabolism (Table 3 and subsequent section).

Taken together, we conclude that gene coexpression pairs are generally robust across multiple network construction scenarios. We suggest that these co-expression resources are highly predictive, reproducible and largely independent of the transcriptional conditions used in their construction.

### Comparing RNA-seq and microarray platforms for network construction

Although most global transcriptional experiments are now conducted using RNA-seq, many organisms have rich resources of publicly available microarray datasets that can be used for co-expression network construction. We therefore compared how our previous *A. niger* microarray co-expression resource compared to the RNA-seq derived network generated in this study. Using an identical Spearman correlation cut-off (0.5), we found that the RNA-seq network had slightly better coverage of the *A. niger* genome when compared to the microarray analysis (9,952 and 9,579, respectively, Table 1), despite having a smaller number of transcriptional conditions used to build the network (128 and 155, respectively). In general, we found limited overlap between RNA-seq and microarray derived sub-networks (Figure 7A), which we speculated could be due to differences in platform sensitivity (32). We thus performed a chemostat bioreactor cultivations with the wildtype strain N402 and the *glaA* over-expression strain B36 (33), an approach that enabled steady-state conditions during growth on the same carbon source (glucose). Extracted RNA was concomitantly analysed by microarray and RNA-seq profiling (see Materials and Methods) for differential gene expression between N402 and strain B36 (Figure 7B and Supplemental File 6). This revealed striking concordance between the two technologies, with 94% and 97% of genes identified as up-regulated and down-regulated by microarray also identified as such by RNA-seq, respectively (Figure 7C).

**Figure 7:**
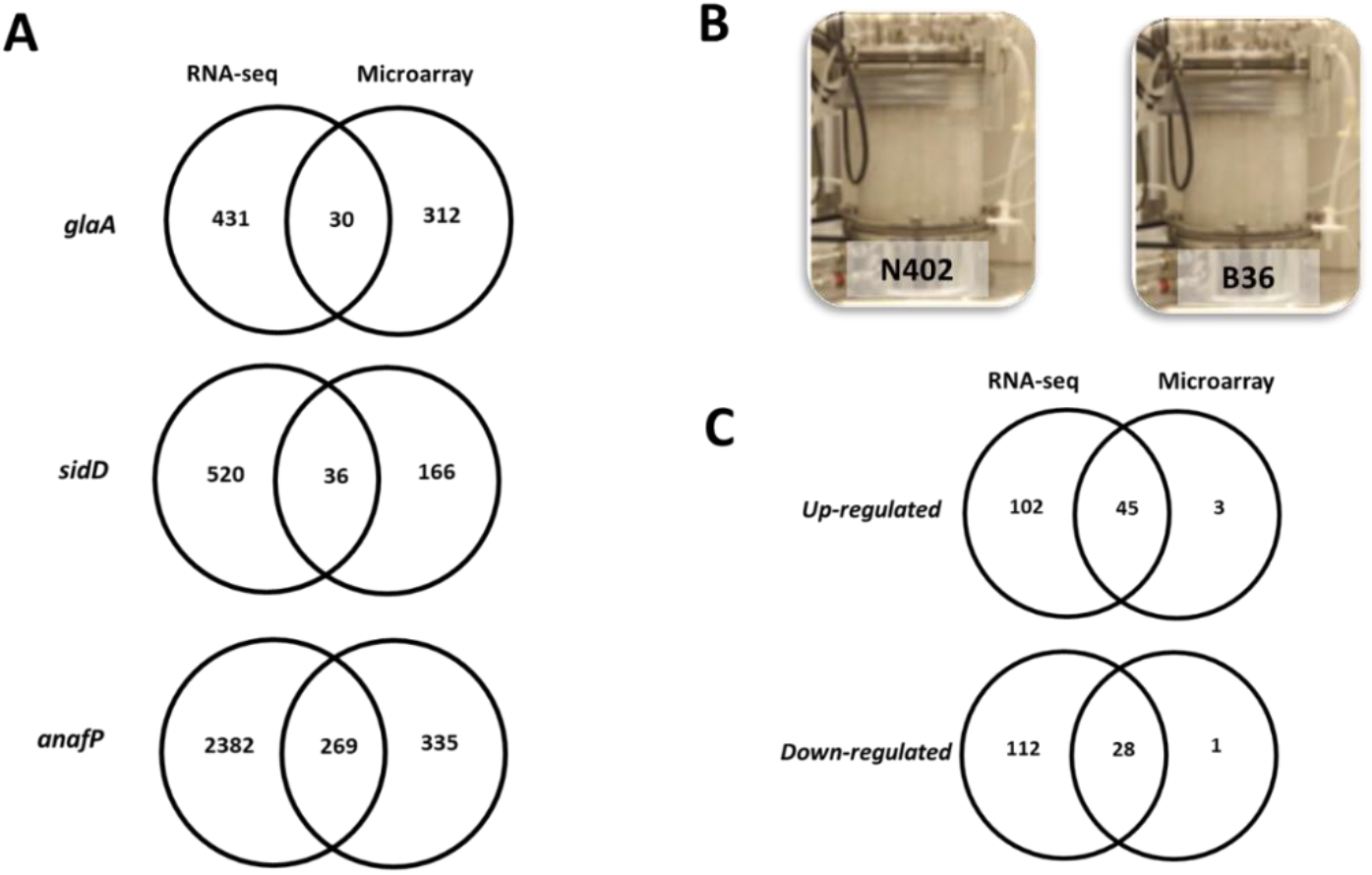
Comparisons between microarray and RNA-seq derived networks. **(A)** Venn diagrams depicting overlap of genes co-expressed with *glaA, sidD*, or *anafP* in RNA-seq (Spearman > 0.25) and microarray (Spearman > 0.5) networks. **(B)** Experimental set up of bioreactors for strain N402 and *glaA* over-expression isolate B36 from which RNA was extracted and microarray/RNA-seq analysis conducted in parallel. **(C)** Overlap between differentially expressed genes identified from RNA-seq and microarray technology.

However, consistent with enhanced sensitivity and specificity of RNA-seq, greater numbers of differentially expressed genes were identified using this approach. We therefore conclude that the differences for the *glaA, sidD* and *anafp* sub-networks (Figure 7A) are mainly explainable by enhanced sensitivity of the RNA-seq profiling technology.

### Putting the RNA-seq co-expression network to the test

Next, we wanted to test the predictive power of a large network generated from the minimum Spearman correlation cut off identified in this study (Table 2). Specifically, the *anafp* sub-networks deduced from the RNA-seq and microarray co-expression resources contained 2,651 and 986 genes, respectively. 2,632 genes were positively correlated (Spearman > 0.25) and 19 negatively correlated (Spearman < −0.31) in the network based on RNA-Seq data (Supplemental File 7), whereas 605 genes were positively correlated (Spearman > 0.5) and 381 genes negatively correlated (Spearman < −0.5) in the microarray-based network (Table 3, (7, 8)). As outlined above and in (7, 8), both networks predict carbon limitation, asexual development, and secondary metabolism to be important processes into which *anafp* gene expression is embedded. However, the network based on RNA-seq data (for GO terms see Table 3) enabled us to zoom into these processes with much greater resolution.

Based on these data, a clear functional enrichment of pectin degrading enzyme encoding genes was observed in the *anafp* positive subnetwork (GO:0045488, p < 0.005). This includes genes predicted to encode polygalacturonases (*pgaA* (An16g06990), *pgaI* (An01g11520), *pgaII* (An15g05370), *pgaE* (An01g14670), *pgxA* (An12g07500), *pgXB* (An03g06740)), two pectin lyases (*pelD* (An19g00270), *pelf* (An15g07160) and others (Supplemental File 7). In order to test the role of *anafp* in *A. niger* pectin metabolism, we generated mutant STS4.10 in which the *anafp* gene was placed under control of the doxycycline titratable Tet-on cassette. Quantitative phenotypic screens on a range of growth media demonstrated various impacts on reduced or overexpression of *anafp*, including, for example, a strong radial growth defect at high temperature when expression was artificially high (Figure 8). Notably, *A. niger* growth on pectin media was clearly impacted by *anafp* expression levels, whereby doxycycline-controlled overexpression caused defective growth on standard minimal media, yet robust growth on pectin (Figure 8). Thus, a novel functional role of AnAFP in pectin utilization was supported by simple growth assays using the STS4.10 strain. In order to generate further evidence that *anafp* is indeed involved in pectin metabolism, we conducted RNA-seq analysis of STS4.10 and VG8.27 control in cultivation in media with 0 or 5 µg/ml Dox (see Materials and Methods), representing null or over expression respectively. Dox induction caused drastic increase in *anafp* expression in STS4.10 (log_2_ 8.07, p < 0.001), and increased or reduced expression of 434 or 407 genes respectively (log_2_ +/-1.5, FDR p < 0.05, Supplemental File 8). Genes with increased expression following *anafp* over expression were enriched in those with a predicted location in the extracellular region (GO0005576, p < 0.00001) and, notably, pectin metabolism (GO:0045488, p < 0.05). Encouragingly, all five differentially expressed pectin genes (*pgaI*, *pgaII*, *abnA*, *rgaeA*, and *abfB*) were also present in the *anafp* co-expression network. Additionally, we found genes with reduced expression following *anafp* over expression were enriched in secondary metabolite processes (GO:0019748, P< 0.001), thus further confirming the predictive power of the large *anafp* co-expression network (Table 3). Taken together, these data provide proof-of-principle that large networks generated from low Spearman thresholds provide robust gene functional predictions and improved resolving power when compared to smaller networks generated from more stringent cut-offs.

**Figure 8:**
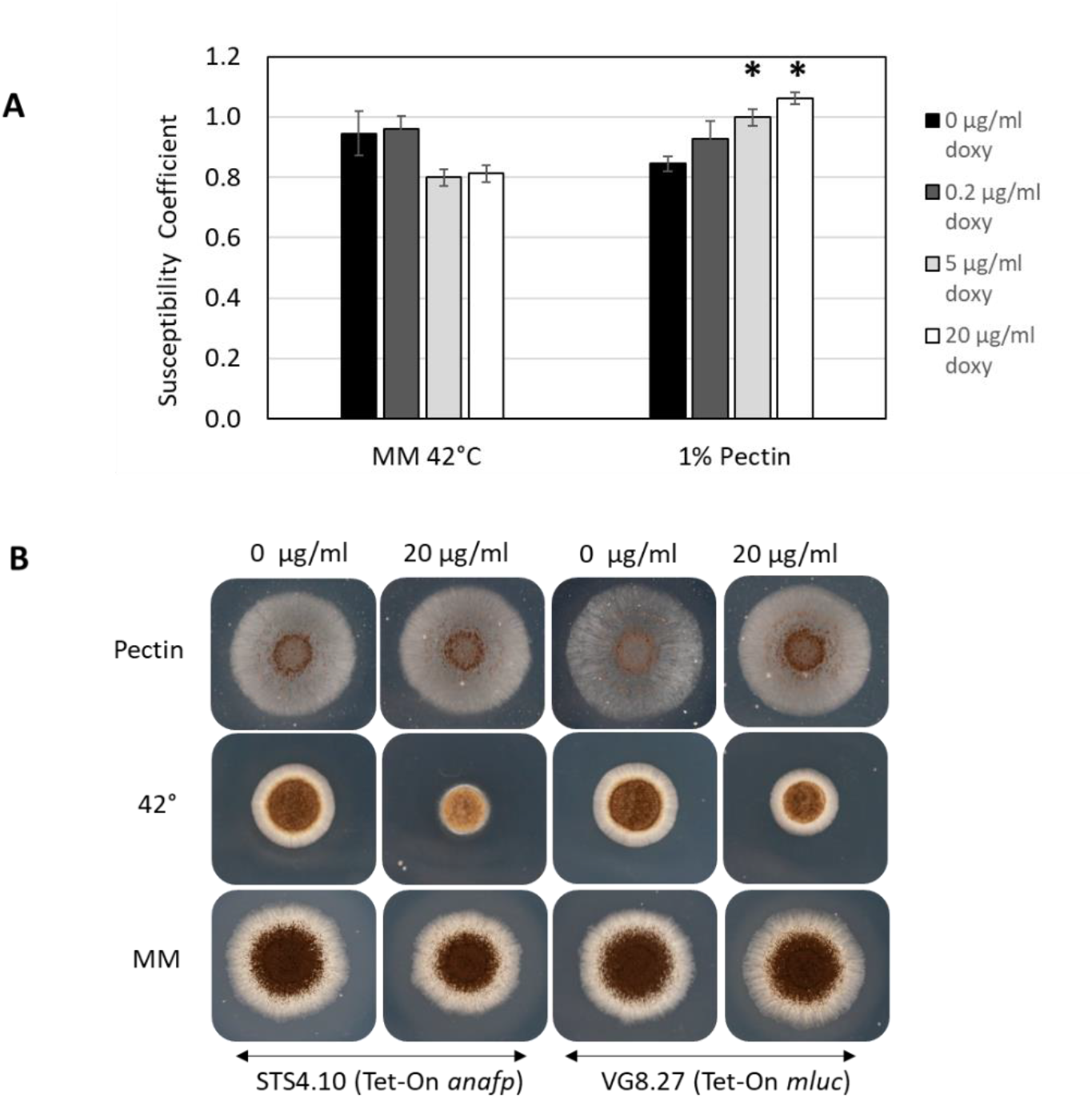
Growth coefficient by colony diameter of *A. niger* STS4.10 (Tet-On *anafp*) relative to control strain VG8.27 (Tet-On *luciferase*). Growth in presence of 0; 0.2; 5 and 20 µg/ml doxycycline, on MM (1% glucose) at 42 °C or on MM with 1% pectin as sole carbon source. n=3, * p-value < 0,05 rel. to 0 µg/ml dox. (B) corresponding colony growth of STS4.10 and VG8.27 at 0 and 20 µg/ml dox, 72 h.

Finally, the *anafp* subnetwork supported the hypothesis that this protein is a major component of nutrient recycling and adaptation of *A. niger* to limited carbon resources (8). If AnAFP does perform such functional role(s), we hypothesized that this protein could be spatially controlled during colony growth, as is the case for various extracellular proteins in *A. niger* and other fungi (29, 34). By placing a luciferase reporter under control of the native *anafp* promoter, we demonstrated that expression occurs in specific colony sections, firstly the colony centre (young colony 18 hours post inoculation), and then immediately behind the advancing hyphal front (maturing colony 36 hours and mature colony 60 hours post inoculation, Figure 9). We noticed expression in condiophores (e.g., at 36 h, Figure 9). Thus, we conclude that the *anafp* promoter is active in carbon-starved vegetative mycelium, present in neighbouring foraging hyphal cells as well as in conidiophores, and still not evenly expressed throughout the mycelium, demonstrating that *anafp* gene expression is under tight spatial and temporal control. These data are consistent with the hypothesis that *anafp* may be an important molecule that connects both nutrient recycling and carbon adaption to ensure survival of the *A. niger* mycelium (29, 34), a notion strongly supported by the co-expression network presented in this study. Taken together, AnAFP may be a nexus that connects crucial biological processes in *A. niger*, and its further functional analysis may enable better understanding of nutrient acquisition, polar growth, cell wall remodelling, protein secretion, mitosis, cell cycle progression and others.

**Figure 9:**
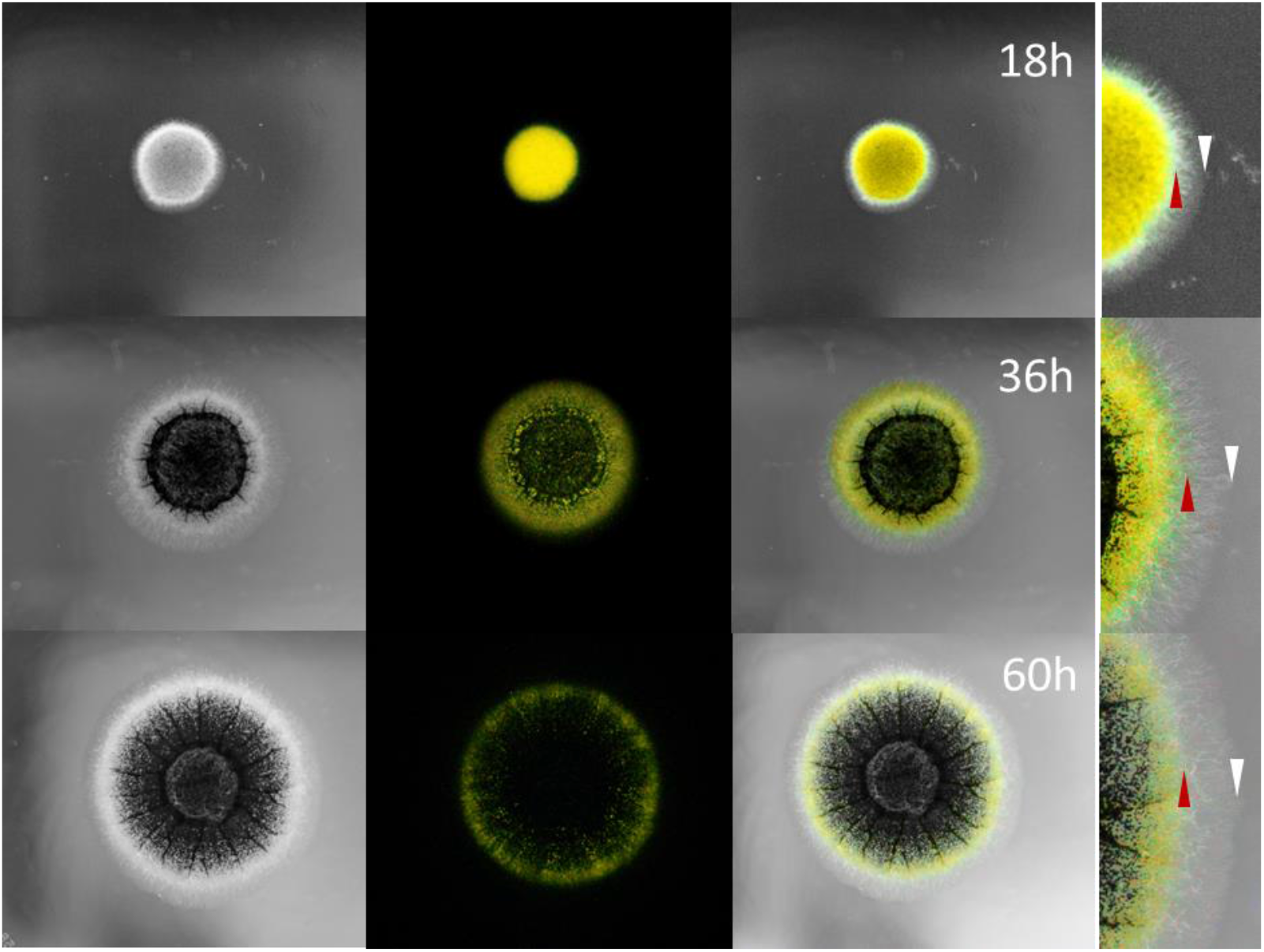
*A. niger* PK2.9 (P*anafp*-*mluc*; Δ*anafp*) long time exposure photography of colony growth at 30°C on CM agar. From left to right: short exposure photography (light), long exposure photography (dark), overlay and corresponding zoom at 18, 36 and 60 h of growth. Green/ yellow luminescence signal depicts the promotor activity of *anafp* via the reporter gene luciferase (*mluc*). Red arrow marks the edge of the luminescence signal. The white arrow marks the edge of the colony.

## Discussion

This study has built and quality control checked an RNA-seq co-expression network for the microbial cell factory *A. niger*. As 70% of predicted genes in this fungus are represented in the network, this resource constitutes a useful update for improving annotation at a near genome level. Encouragingly, this resource will help address the hypothetical protein problem, whereby an estimated 50% of genes lacking any functional prediction can now be interrogated based on the transcriptional network in which they are embedded. We have made these data fully available at FungiDB, where users can interrogate gene sub-networks using a comprehensive analysis pipeline (e.g., searching gene sub-networks for enriched GO terms, metabolic pathways, secreted proteins, etc.).

We have also tested and developed guidelines for several aspects of network construction. Arguably, the most important discovery is confirmation that an informative, near genome-wide co-expression network can be constructed from 32 expression datasets. Consequently, high quality co-expression networks can be made for many fungi from readily available RNA-seq data, including human infecting, plant infecting, industrial cell factories or model organisms. One interesting aspect from our study was the limited impact of RNA-seq input on gene sub-network content, either by randomly selecting different networks, or purposely selecting biased datasets (e.g., without nitrogen limitation). Our analysis is therefore consistent with previous studies which show ‘safety in numbers’, i.e., provided a minimum threshold of RNA-seq experiments are included, the network will be of sufficient quality irrespective of the transcriptional conditions which are used (35, 36). This is especially encouraging for fungi in which *in vitro* growth is either challenging or yet to be established, e.g. for emerging plant or human infecting pathogens.

Our study also advises minimum Spearman correlation coefficients for various amounts of data input. As expected, lower thresholds can be used for networks generated from more data (e.g., 0.25 and 0.56 for 128 and 32 experiments, respectively). Obviously, the precise threshold used to define co-expression pairs will depend on the end user-e.g., for time and reagent intensive mutant construction for mechanistically proving gene function through wet-lab experiments, much higher thresholds than our advised minimum should be used. However, for predicting gene function with too stringent Spearman thresholds in the gene sub-networks, we argue that hypothesis generation might not be comprehensive enough.

Another observation in this study was that RNA-seq and microarray derived gene sub-networks differ. We argue that this is explainable due the differing sensitivity of the technologies as also discussed by others (32) and due to variations in applied Spearman thresholds that one can use. However, due to the ‘safety in numbers’ principle, and the fact that the networks built here and in our previous microarray study were made from over 100 conditions, we consider it unlikely that variation for experimental conditions effected the networks. Importantly, by concomitantly profiling identical RNA using microarray and RNA-seq, and confirming remarkable concordance between the two platforms, we rule out the hypothesis that transcriptional experiments from older technology, i.e., microarray, should be avoided due to poor calls in differential transcript abundance. Therefore, the many thousands of publicly available gene expression datasets that have been made with microarrays can still be used with confidence for network construction.

One notable observation from the RNA-seq co-expression datasets generated in this study was the limited number of negative correlations (∼517,000) when compared to positive correlation (∼5 million, Table 1). We hypothesize that this probably reflects the fact that most processes and pathways require expression of multiple genes, thus making positive correlations more frequent. Another possible explanation is that derepression of functionally related gene cohorts tends to cause positive co-expression, for example during carbon catabolite derepression (37). While it is extremely difficult to estimate the frequency of gene repression/derepression in any organism, this certainly constitutes a major regulatory mechanism, occurring at the level of epigenetics (i.e., via chromatin structure), transcription (e.g., repressor elements binding to promoter regions), and RNA interference. Thus, we consider the low numbers of negative correlations to be an interesting observation that may be substantiated by construction of other networks in different organisms.

Finally, functional analysis of one gene of our interest, *anafp,* by titratable expression confirmed new leads for understanding the connection of global regulation phenomena driving *anafp* expression with notably pectin metabolism. Improved non-intuitive predictive power from the co-expression networks generated in this study is facilitated by higher sensitivity of RNA-seq when compared to microarray derived networks, and, additionally, the use of low correlation thresholds to build large gene subnetworks.

## FUNDING

The work was funded by the Deutsche Forschungsgemeinschaft (DFG, GRK2473 ‘Bioactive Peptides’—project number 392923329; DFG project AnAFP ME2041/11-1; DFG project MorphAN ME2041/13-1; DFG project SPP 1934 DispBiotech ME 2041/5-2; DFG project Interzell ME2041/12-1).

---

**Supplemental Figure 1:**
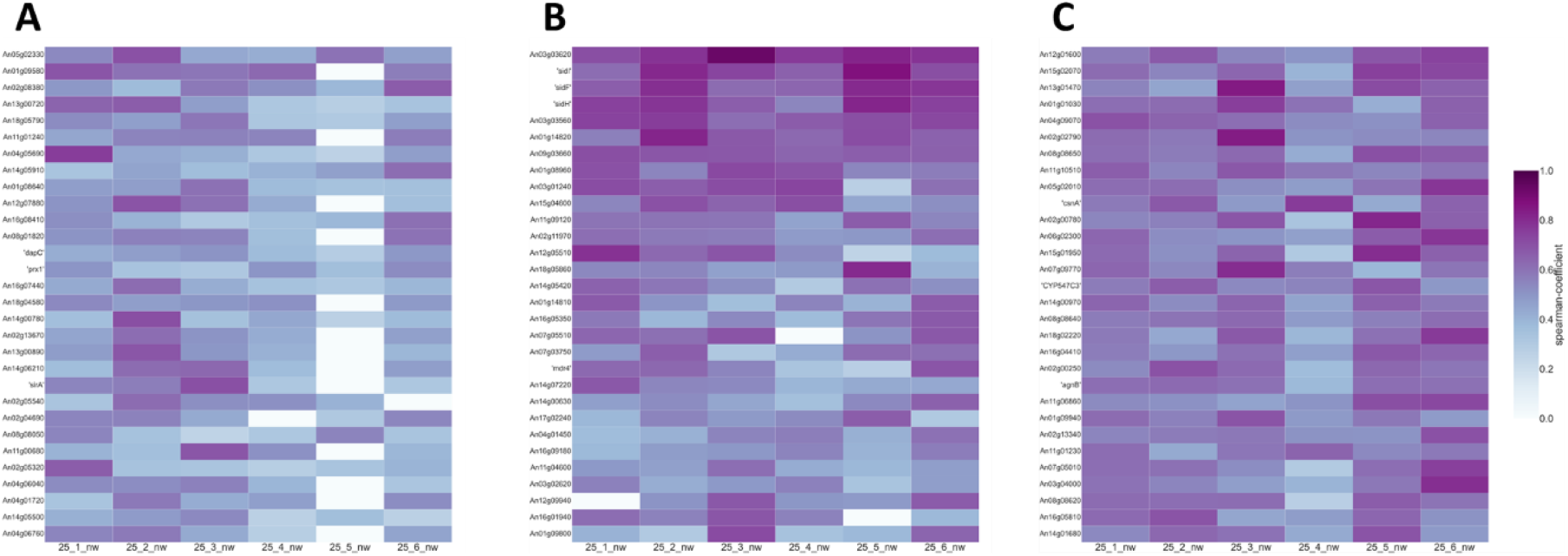
Correlation coefficients for *glaA, sidD*, and *anafP* across six different test networks derived from 32 experimental conditions that were randomly chosen. Top 30 highest co-expressed genes are depicted for either *glaA* (A), *sidD* (B), or *anafP* (C). Each column represents a distinct network derived from 64 randomly selected RNA-seq experiments. Colour intensity correlates with Spearman correlation coefficient.

## Notes

### Competing Interest Statement

The authors have declared no competing interest.

